# Liquid-liquid phase separation mediated immune evasion of RSV against OAS-RNase L pathway

**DOI:** 10.1101/2025.06.24.661296

**Authors:** Woo Yeon Hwang, Michael G. Rosenfeld, Soohwan Oh, Young-Chan Kwon

## Abstract

Respiratory syncytial virus (RSV) infection is the major cause of severe respiratory illnesses in infants and older adults. RSV forms phase-separated biomolecular condensates called inclusion bodies (IBs), which serve as hubs for viral replication. However, the contribution of IBs to host immune response evasion remains elusive. We report that RSV IBs protect viral RNA from the 2′-5′ oligoadenylate synthetase (OAS)-RNase L pathway, a critical antiviral defense mechanism that cleaves viral and cellular RNAs. RSV infection did not activate the OAS-RNase L pathway, and ectopically activated RNase L did not suppress viral replication. In RSV-infected cells, double-stranded RNA (dsRNA) was efficiently sequestered within liquid–liquid phase separation (LLPS)-mediated IBs, rendering its detection challenging. LLPS perturbation caused dsRNA release from IBs into the cytosol. dsRNA extracted from infected cells, which lacked LLPS shielding, triggered OAS-RNase L pathway activation. Thus, LLPS-driven IBs structurally sequester viral RNA, facilitating RSV to evade RNase-dependent genomic RNA degradation mediated by the OAS-RNase L antiviral pathway.

## Introduction

Membrane-less organelles (MLOs) such as inclusion bodies (IBs) and nuclear bodies are dynamic cellular compartments present in different cell regions. These structures are formed via liquid–liquid phase separation (LLPS) of biomolecules, RNAs, and proteins with intrinsically disordered regions(Hyman *et al*, 2014; Protter *et al*, 2018; Lichtinger *et al*, 2021). Unlike traditional organelles, MLOs lack physical barriers, facilitating the reversible and dynamic assembly of distinct liquid-like droplets that can be visualized under a microscope(Nott *et al*, 2016; Li *et al*, 2022; Gomes & Shorter, 2019). This unique property is exploited by viruses to improve replication efficiency, evade host immunity, and remodel cellular environments, making them the main therapeutic targets for antiviral strategies(York, 2021).

Respiratory syncytial virus (RSV) is a major cause of acute respiratory tract infections worldwide, with most people being exposed to the virus by the age of two(Chanock *et al*, 1957; Glezen *et al*, 1986). While RSV typically causes mild symptoms resembling those of the common cold, it can lead to severe diseases such as bronchiolitis and pneumonia in vulnerable populations, including infants, older adults, and immunocompromised individuals(Falsey *et al*, 2005; Peebles & Graham, 2005). Effective vaccines for direct administration to infants remain unavailable, despite recent advancements in RSV vaccine development, including regulatory approvals for vaccines aimed at older adults and maternal immunization strategies(Fukushima *et al*, 2023; Cardona *et al*, 2025; Wilson *et al*, 2023; Papi *et al*, 2023; Dieussaert *et al*, 2024; Schmoele-Thoma *et al*, 2022; Battles & McLellan, 2019). RSV is an enveloped, negative-sense, single-stranded RNA virus belonging to the *Orthopneumovirus* genus of the *Pneumoviridae* family(Afonso *et al*, 2016). It has a non-segmented RNA genome of approximately 15 kb in length and encodes 10 genes that are translated into 11 proteins. Among these viral proteins, nucleoprotein (N), phosphoprotein (P), polymerase (L), and the transcription factor M2-1 accumulate in the IBs (MLO subtype) that form in the cytoplasm of RSV-infected cells(Norrby *et al*, 1970; Wileman, 2007; Carromeu *et al*, 2007; García-Barreno *et al*, 1996; Santangelo & Bao, 2007). LLPS generates these IBs, which serve as viral replication hubs, concentrating viral genomic RNA and core replication machinery(Li *et al*, 2022; Rincheval *et al*, 2017; Galloux *et al*, 2020; Zhang *et al*, 2024). Notably, LLPS hardening blocks RSV replication, underscoring the functional criticality of phase separation in the viral life cycle(Risso-Ballester *et al*, 2021). While IBs are well-recognized as the viral factories supporting RNA synthesis, emerging evidence suggests that they may also contribute to immune evasion, possibly by leveraging their dynamic properties to compartmentalize viral RNA and shield it from host defenses(Lifland *et al*, 2012; Jobe *et al*, 2020). However, these additional functions are yet to be completely elucidated.

Double-stranded RNA (dsRNA) is an inevitable byproduct of viral replication, particularly RNA viruses such as RSV. Almost all organisms exhibit pathogen recognition receptors capable of detecting dsRNAs and initiating antiviral immune responses(Chen & Hur, 2022). The 2′-5′ oligoadenylate synthetase (OAS) system induced by the interferon (IFN) signaling pathway is one such sensor that detects dsRNA. OASs become catalytically active upon binding to dsRNA and synthesize 2′-5′ oligoadenylates (2-5A) from ATP molecules, acting as secondary messengers to activate ribonuclease L (RNase L). Activated RNase L dimerizes and cleaves cellular and viral single-stranded RNA, resulting in the indiscriminate suppression of protein expression from viral and host sources(Kristiansen *et al*, 2011; Silverman, 2007). The OAS-RNase L pathway has been confirmed to exert antiviral activity against different viruses, including the Sindbis virus, West Nile virus, dengue virus, and SARS-CoV-2(Li *et al*, 2016, 2021; Lin *et al*, 2009; Kwon *et al*, 2013), whereas some viruses, such as influenza and Zika viruses, use diverse strategies to evade this antiviral mechanism(Whelan *et al*, 2019; Zhao *et al*, 2012; Min & Krug, 2006; Drappier & Michiels, 2015). However, the antiviral effects of the OAS-RNase L pathway on RSV have not yet been clearly established.

Herein, we investigated the mechanism by which RSV evades the antiviral activity of the OAS-RNase L pathway. By measuring viral RNA and ribosomal RNA (rRNA) cleavage, we found that RSV efficiently circumvents the host dsRNA-sensing mechanism, including OAS activation, by forming IBs driven by LLPS. Using dsRNA-specific antibodies, we observed that viral RNA was sequestered within the IBs, protecting them from recognition and degradation by activated RNase L. These findings show that LLPS-driven IBs serve as replication hubs and protective compartments, facilitating RSV to escape RNase-dependent degradation of genomic RNA mediated by the OAS-RNase L antiviral pathway.

## Results

### RSV infection does not activate the OAS-RNase L pathway

The OAS-RNase L pathway is activated against infections from a diverse range of viruses(Li *et al*, 2016, 2021; Lin *et al*, 2009; Kwon *et al*, 2013). We infected the wild-type A549 cells with RSV and assessed RNase L activation via rRNA cleavage, the hallmark of RNase L activation, at various times after infection to investigate whether RSV infection activates the OAS-RNase L pathway. We used the prototypic strains RSV A2 and RSV 18537, which represent historically established reference strains for the RSV subgroups A and B. We transfected cells with polyinosinic:polycytidylic acid (poly (I:C)) as a surrogate for dsRNA to activate the OAS-RNase L pathway as a positive control(Li *et al*, 2016). Despite efficient viral replication over time, rRNA cleavage was not observed in the cells infected with either RSV strain, regardless of the time point (Fig 1A and B). Also, rRNA cleavage was not observed in the HEp-2 cells, which are highly susceptible to RSV infection (Fig EV1). Contrastingly, rRNA cleavage was detected in Zika virus-infected cells, as previously reported (Fig EV2)(Whelan *et al*, 2019). Furthermore, RSV infection with different multiplicities of infection (MOI), ranging from 0.1–5, did not induce rRNA cleavage (Fig 1C). Next, we determined the protein and mRNA expression levels of genes related to the OAS-RNase L pathway in the RSV-infected cells (Fig 1D and E). As expected, *OAS1*, *OAS2*, and *OAS3*, which are the components of interferon-stimulated genes, were upregulated after RSV infection. Thus, RSV infection did not activate the OAS-RNase L pathway despite the upregulation of the expression levels of its components.

**Figure 1-.**
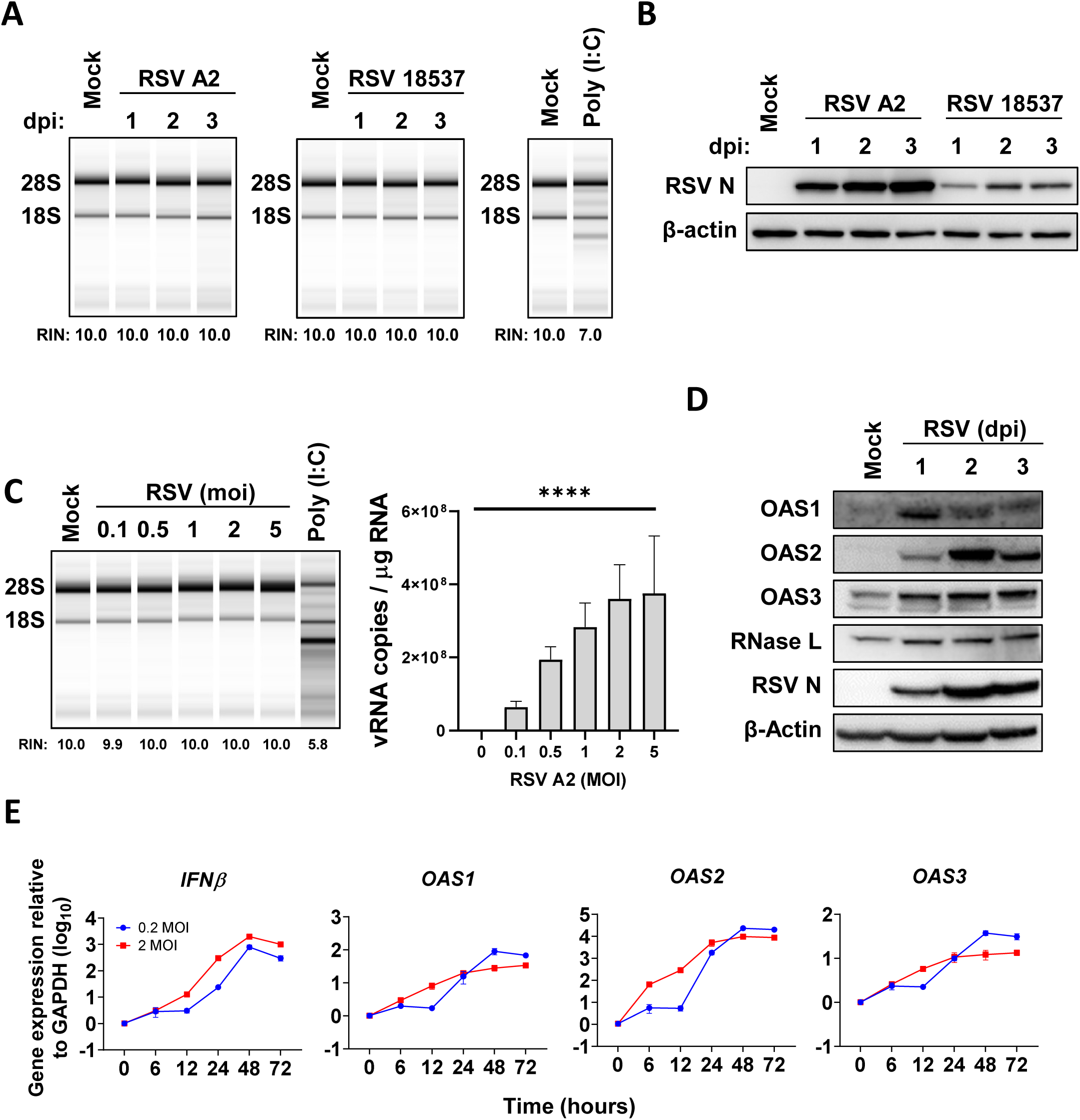
Respiratory syncytial virus (RSV) infection does not elicit oligoadenylate synthetase (OAS)-RNase L activation. (A) A549 cells were infected with RSV A2 or RSV 18537 at an MOI of 2. Cells were collected at indicated time points. rRNA cleavage and RNA integrity number (RIN) were analyzed using an RNA TapeStation System. (B) Protein levels were examined using immunoblotting with anti-RSV N and anti-β-actin antibodies. (C) A549 cells were infected with RSV A2 at the indicated multiplicities of infection (MOIs) and lysed 2 d after infection. Viral RNA levels were quantified using quantitative reverse transcriptase-polymerase chain reaction (RT-qPCR). (D) After RSV infection at an MOI of 2, protein levels of the OAS-RNase L pathway components (OAS1, OAS2, OAS3, and RNase L) were analyzed at the indicated time points via immunoblotting. (E) mRNA levels related to the IFN-β and OAS-RNase L pathway in the RSV-infected cells were measured using RT-qPCR and normalized to GAPDH. Data represent mean ± standard errors of the mean from three independent experiments. Statistical significance was determined via one-way analysis of variance (C); ****P < 0.0001.

### RSV replication persists despite RNase L activation

Ectopically activated RNase L suppresses viral replication, even in the absence of OAS-RNase L pathway activation via viral infection.(Whelan *et al*, 2019; Park *et al*, 2014). Thus, we aimed to examine the effect of ectopically activated RNase L on RSV replication since RSV infection does not activate the OAS-RNase L pathway. Cells were infected with RSV for 24 h and subsequently transfected with poly (I:C) to activate RNase L. rRNA cleavage was observed in all poly (I:C)-transfected cells, regardless of RSV infection (Fig 2A). These observations showed that RSV does not impede the OAS-RNase L pathway activated by ectopic dsRNA. Notably, RNase L, activated by different poly (I:C) doses, did not inhibit RSV replication (Fig 2B).

**Figure 2-.**
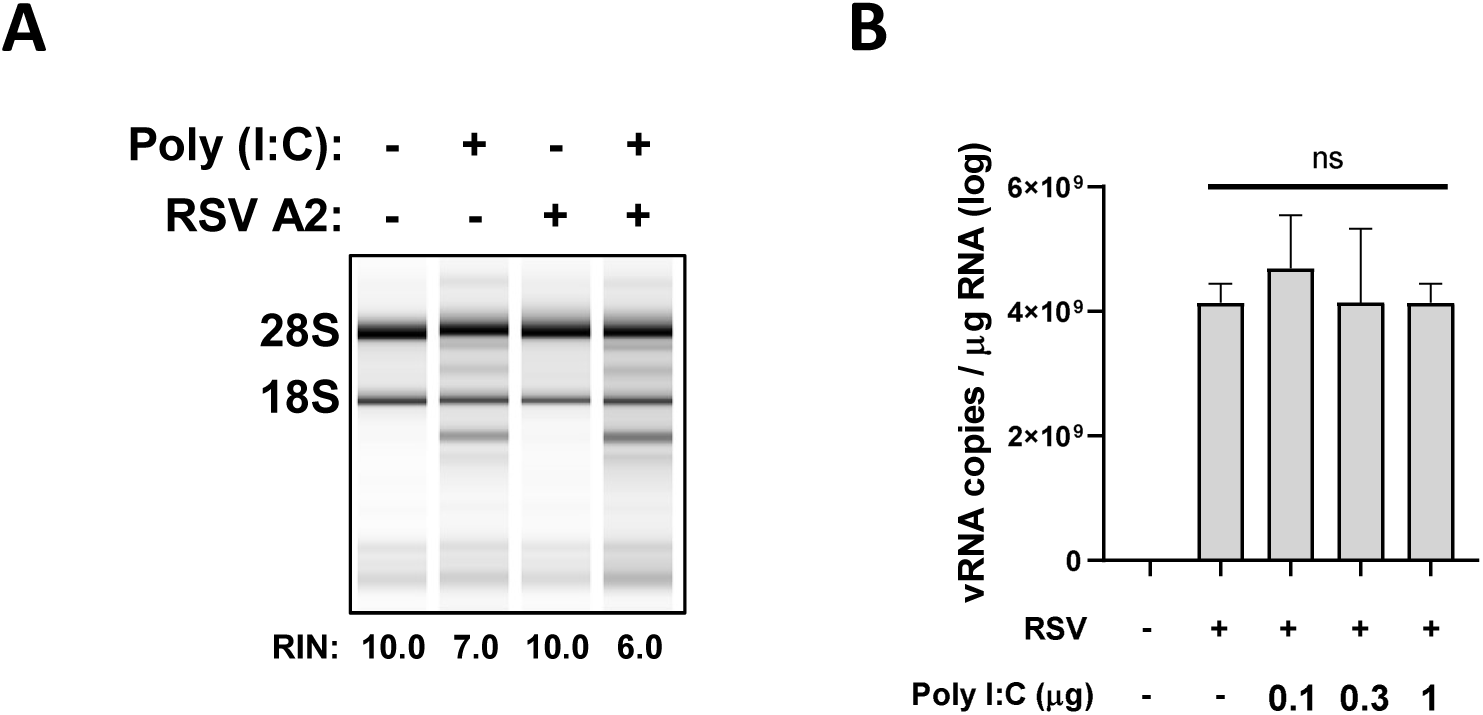
Activated RNase L does not impair RSV replication. (A, B) A549 cells infected with RSV A2 (MOI = 2) were transfected with (A) 1 μg or (B) indicated amount of poly(I:C) at 24 h post-infection. RNA was isolated 24 h later and analyzed for rRNA cleavage and viral RNA levels. Statistical significance was determined using a one-way analysis of variance (B); ns: not significant. Data represent mean ± standard errors of the mean.

The distinct set of OAS isoforms generated by alternative splicing show different intensities of antiviral activity against diverse viruses(Li *et al*, 2016; Lin *et al*, 2009). Moreover, the A549 cells used in this study did not produce the OAS1 p46 isoform, as demonstrated in our previous study(Kwon *et al*, 2013). Different OAS isoforms, including OAS1 p46, were overexpressed, followed by RSV infection to enhance the A549 cells’ responsiveness to dsRNA. However, rRNA cleavage was not observed (Fig EV3). Overall, these results indicate that RSV viral RNA generated during replication evades OAS-mediated dsRNA-sensing and is likely protected from degradation by the activated OAS-RNase L pathway.

### Structural masking of viral dsRNA underlies RSV’s evasion of cytosolic sensing

During virial replication, dsRNA is generated, although RSV infection does not activate dsRNA-dependent OASs. dsRNA and viral protein (F, RSV fusion protein) were probed using specific antibodies in an immunofluorescence assay (IFA) in RSV-infected cells for 48 h to evaluate the extent of dsRNA production during RSV replication (Fig 3A). Surprisingly, dsRNA was not detected, although the viruses efficiently replicated, as indicated by viral protein (F) expression (Fig 3B). Contrastingly, ZIKV-infected cells showed clear dsRNA signals along the viral protein (E, ZIKV envelope protein). Because dsRNA is inevitably produced during the RNA virus replication process and is a known target for the host defense system, considering its absence in virus-infected cells is illogical(Chen & Hur, 2022). We resolved this contradiction by conducting an experiment involving immunoblotting to probe for dsRNA in total RNA samples isolated from RSV-infected cells using a previously reported methodology (Fig 3A)(de Faria *et al*, 2023). The dsRNA signal was detected in the RSV-infected samples (Fig 3C), and was further confirmed using another anti-dsRNA antibody (9D5) targeting a different epitope (Fig EV4). These results suggest that the dsRNA produced during RSV replication is structurally obscured, preventing it from being sensed by OASs.

**Figure 3-.**
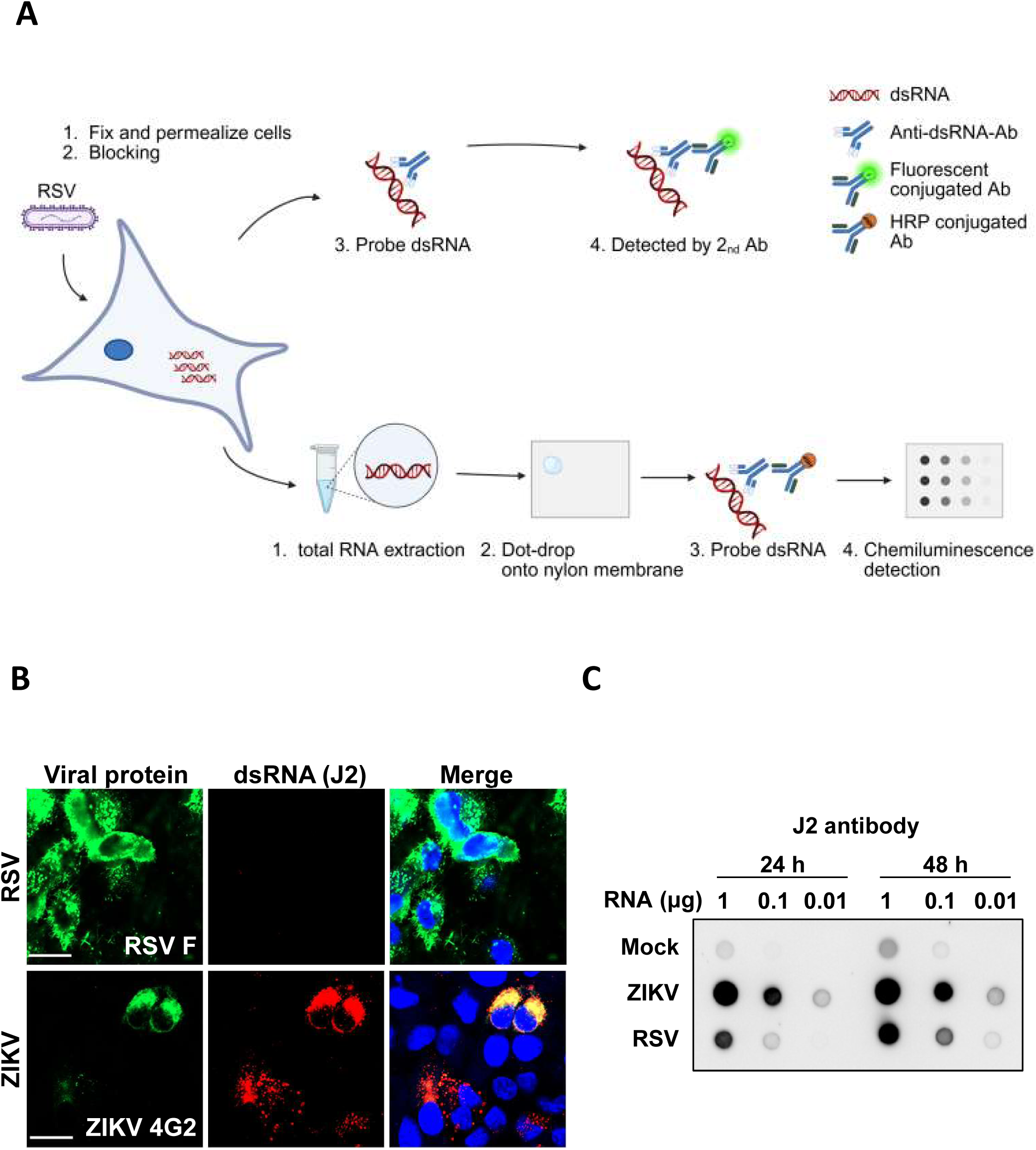
dsRNA generated during RSV replication is present but remains undetectable via immunofluorescence assay (IFA). (A) overview about detecting dsRNA through IFA and dot-bloting. (B) A549 cells infected with RSV A2 (MOI = 2) were fixed at 2 dpi and stained with anti-dsRNA (J2, red) and anti-RSV F antibodies (green). Images were acquired using an LSM 980 confocal microscope. ZIKV-infected cells (MOI = 2) stained with anti-ZIKV envelope antibody (green). (C) Total RNA from mock-, RSV-, or ZIKV-infected cells was transferred to a nylon membrane. dsRNA was detected using anti-dsRNA antibody (J2). Scale bar: 20 μm.

### IBs sequester viral dsRNA

During infection, RSV induces cytoplasmic inclusion formation to produce IBs(Norrby *et al*, 1970; Wileman, 2007). These results were confirmed by staining with an anti-N antibody (Fig 4A), which revealed detectable IBs at 12 h post-infection, with progressive expansion over time. IBs were enriched with viral proteins involved in viral replication, namely, N, P, and M2-1 (Fig 4B), as recently reported. L protein, an RNA-dependent RNA polymerase of RSV, is also known to localize within IBs(Carromeu *et al*, 2007; García-Barreno *et al*, 1996; Santangelo & Bao, 2007). Additionally, a previous study showed that newly synthesized viral RNA is situated within IBs(Rincheval *et al*, 2017). These data confirmed that IBs were located within the newly synthesized viral RNA.

**Figure 4-.**
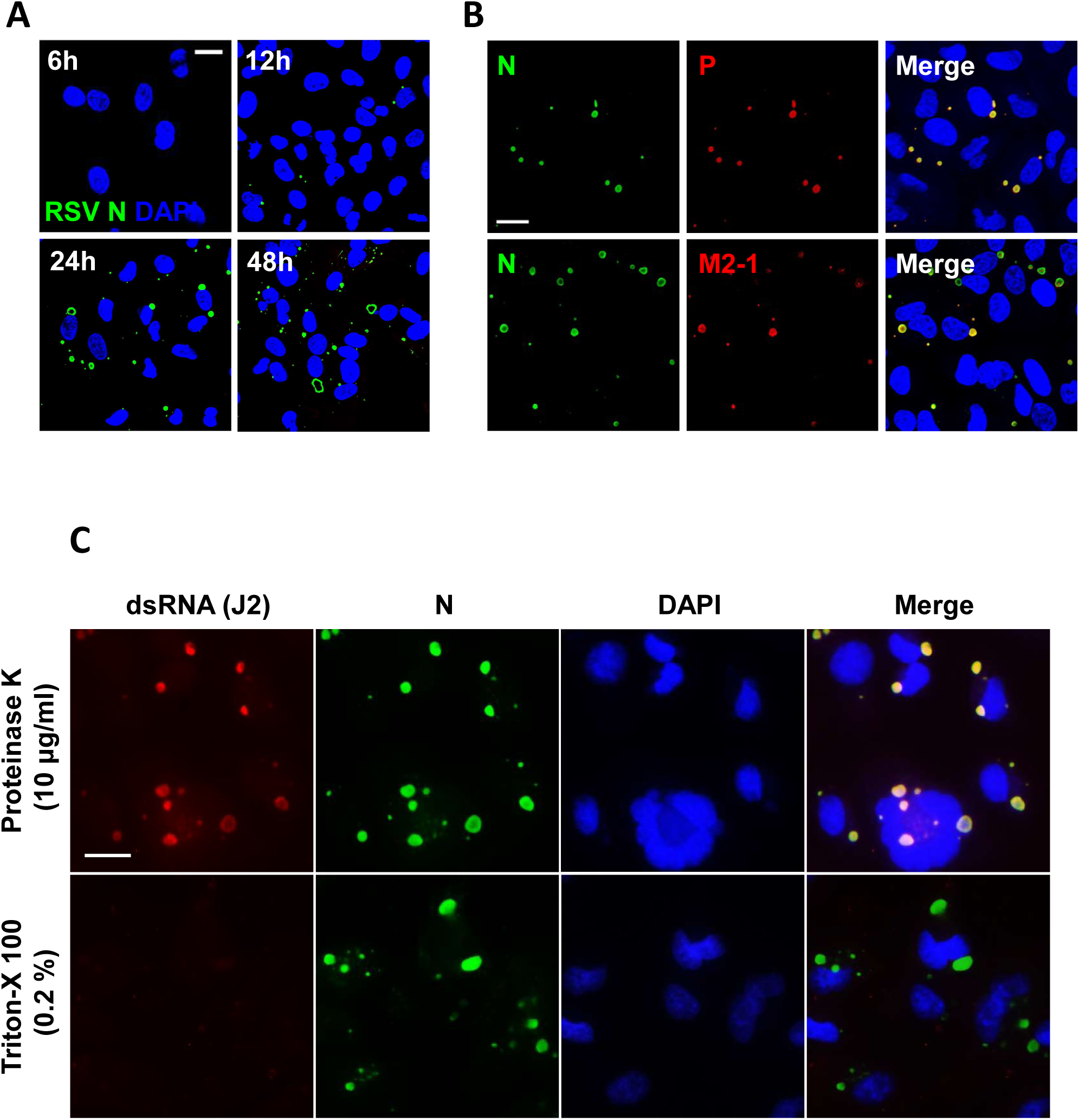
Sequestering dsRNA within inclusion bodies (IBs), which are enriched with viral replication factors. (A, B) A549 cells infected with RSV A2 (MOI = 2) were stained with anti-N (green), anti-P (red), and anti-M2-1 (red) antibodies, as well as 4’,6-diamidino-2-phenylindole (blue) at (A) indicated time points or (B) 24 h. Images were acquired using an LSM 980 confocal microscope. (C) RSV-infected cells at 24 h post-infection were treated with proteinase K before staining. Images were obtained using a Nuance FX Multispectrum Imaging System. Scale bar: 20 μm.

Stained N or P proteins, identified as the minimal components necessary for IB formation, were commonly observed to be unevenly distributed within the IBs, which appeared hollow, as repeatedly reported(Rincheval *et al*, 2017; Galloux *et al*, 2020). This was especially evident 24 h post-infection (Fig 4A and B). We hypothesized that IBs hinder antibody accessibility to dsRNAs and demonstrated this by treating cells with proteinase K during permeabilization, which is commonly used in fluorescence in situ hybridization assays to remove RNA-binding proteins(Zimmerman *et al*, 2013). Notably, proteinase K treatment allowed dsRNA detection in the RSV-infected cells, where it co-localized with the RSV N protein within the IBs (Fig 4C). Visual evidence clearly showed that dsRNA was obscured by the IBs, which might explain why RSV dsRNA was detected only in the lysates and not in the IFA (Fig 3). Collectively, our results suggest that IBs function as viral replication factories and sequester dsRNAs as by-products. Furthermore, this sequestration may prevent OASs from sensing dsRNAs during RSV replication.

### Suppression of IBs induces dsRNA leakage

The morphology and properties of IBs formed during RSV replication suggest they are condensates generated through LLPS(Galloux *et al*, 2020; Zhang *et al*, 2024). To disrupt these structures, RSV-infected cells were treated with 5% 1,6-hexanediol (1,6-HD), a commonly used agent that dissolves LLPS assemblies(Li *et al*, 2022; Geiger *et al*, 2021). The cells were subsequently fixed at the indicated time points and stained. Over time, dsRNA was released from the IBs into the cytosol after 1,6-HD treatment (Fig 5A). This observation was further confirmed using another anti-dsRNA antibody (9D5) co-stained with P protein (Fig EV5). A portion of the dsRNA was observed at the periphery of the IBs in the early stages following 1,6-HD exposure for 1 min (Fig 5B). After 30 min from exposure, N protein was also released into the cytosol; however, this was not observed for P protein. This finding reinforces the notion that RSV IBs use a form of LLPS to sequester dsRNA, thereby protecting it.

**Figure 5-.**
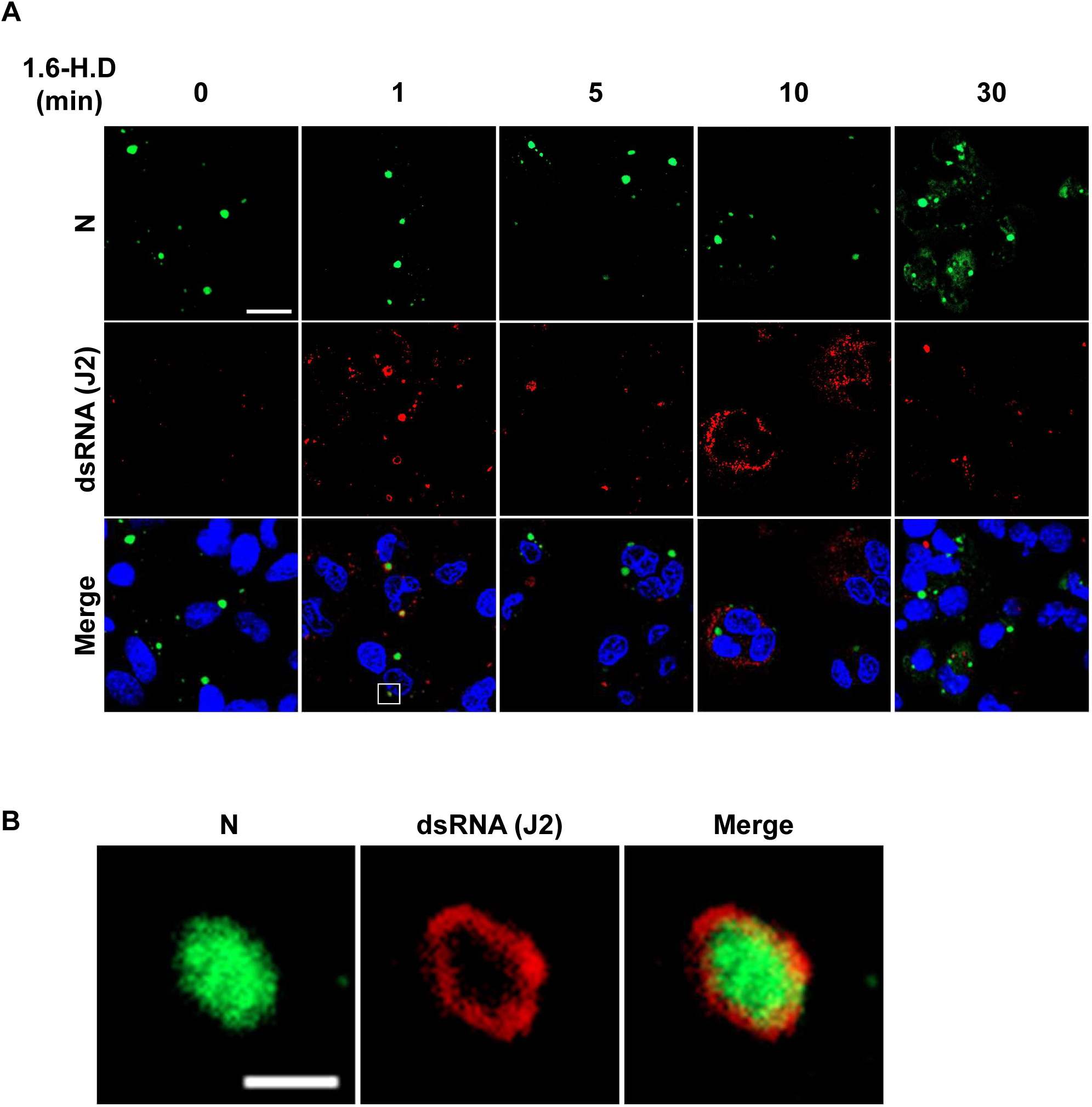
IB disruption with a liquid–liquid phase separation (LLPS) inhibitor induces dsRNA leakage. (A, B) A549 cells were infected with RSV A2 at a MOI of 2 for 24 h. Before cells were fixed, 5% 1,6-hexanediol (1,6-HD) was treated at the indicated durations. Cells were stained with anti-N antibody (green) and anti-dsRNA (J2, red); scale bar: 20 μm. Magnified views of the boxed areas are illustrated (B); scale bar: 2 μm.

### Naked dsRNA isolated from RSV-infected cells induces OAS-RNase L pathway activation

We isolated naked dsRNA from RSV-infected cells by performing a pull-down assay using a dsRNA-specific antibody, as previously reported (Fig 6A), to investigate the dsRNA-sequestering function of IBs(Decker *et al*, 2019). Isolated dsRNA was confirmed via dot blot assay using anti-dsRNA antibodies (Fig 6B; Fig EV6A). Subsequently, we observed rRNA cleavage after the transfection of purified, naked dsRNA from RSV-infected cells (Fig 6C). This trend was consistently observed when total RNA from RSV- or mock-infected cells was used for transfection (Fig EV7). Thus, dsRNA, generated during RSV replication, is protected by IBs, even though it is detectable via the OAS-RNase L pathway.

**Figure 6-.**
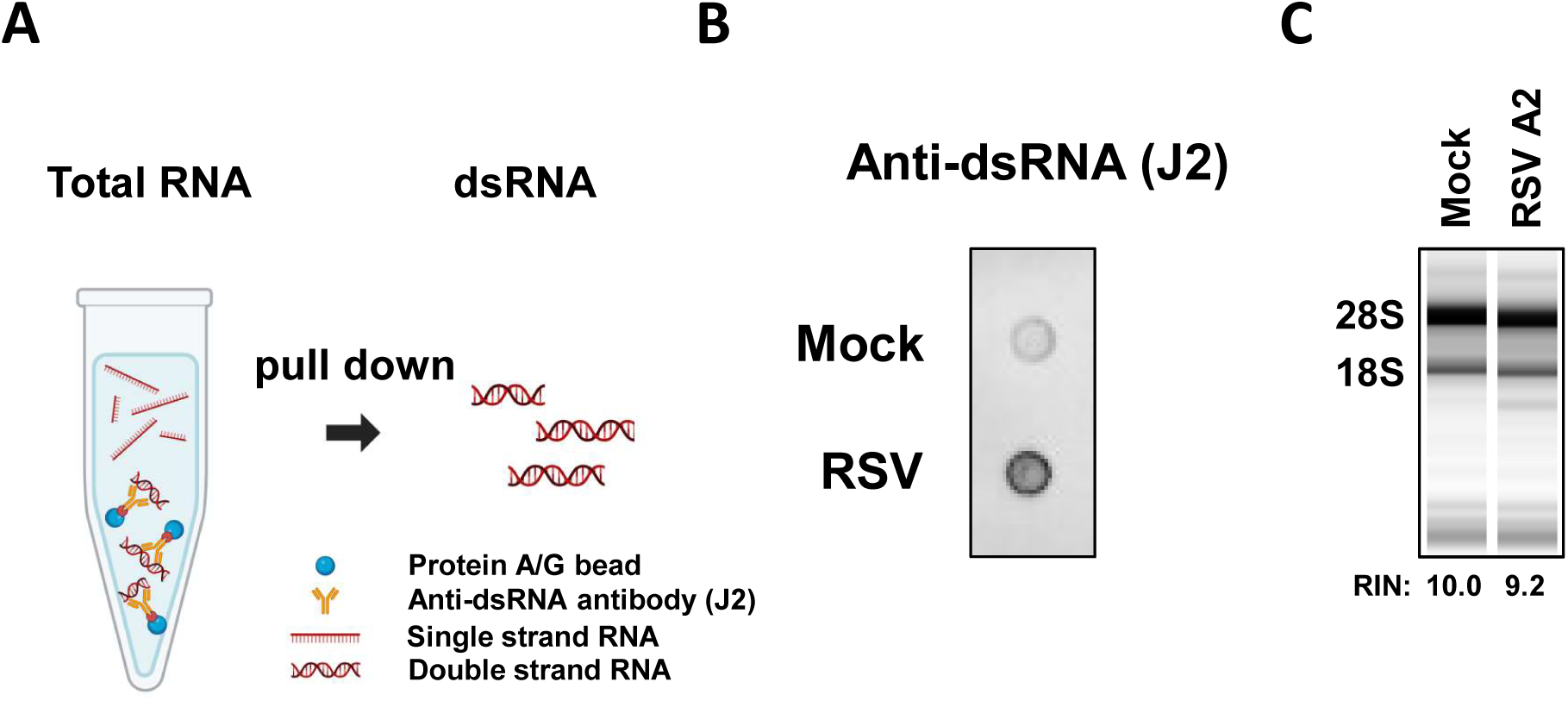
Naked dsRNA isolated from RSV-infected cells induces OAS-RNase L activation. (A) Total RNA extracted from the RSV- or mock-infected cells was used in a pull-down assay with anti-dsRNA antibody pre-bounded to protein A/G beads. (B) dsRNA enrichment was confirmed via dot blotting using an anti-dsRNA antibody (J2). (C) After the purified dsRNA was transfected into the A549 cells, total RNA harvested at 1 d post-transfection was analyzed using the RNA TapeStation System.

## Discussion

The OAS-RNase L pathway, an IFN-stimulated antiviral mechanism, is crucial for the innate immune defense against viral infections(Kristiansen *et al*, 2011; Silverman, 2007). This pathway is initiated following the detection of dsRNA produced during RNA virus replication via OASs. Additionally, activated RNase L indiscriminately degrades viral and host RNA, suppressing viral replication(Li *et al*, 2016, 2021; Lin *et al*, 2009; Kwon *et al*, 2013). Contrastingly, viruses use diverse strategies to evade this pathway(Whelan *et al*, 2019; Min & Krug, 2006; Drappier & Michiels, 2015; Silverman & Weiss, 2014). The relationship between RSV infection and the OAS-RNase L pathway has been reported in several studies(Barnard *et al*, 1999; Player *et al*, 1998; Cirino *et al*, 1997; Leaman *et al*, 2002). However, the precise molecular mechanisms involved remain to be completely elucidated. Herein, we showed that RSV evades the OAS-RNase L pathway by sequestering viral RNA within LLPS-driven IBs, preventing OASs from detecting dsRNA. Consequently, even ectopically activated RNase L fails to suppress RSV replication.

Unexpectedly, RSV infection did not activate the OAS-RNase L pathway, even though infection upregulated the expression levels of OAS-RNase L pathway components. Furthermore, attempts to increase OASs’ sensitivity by overexpressing OAS isoforms did not result in pathway activation. Additionally, RSV did not halt the ectopic activation of the OAS-RNase L pathway but this activation still failed to inhibit RSV replication. Numerous studies have discovered diverse strategies for viruses to evade OAS-RNase L, such as generating viral genome reservoirs (e.g., Zika virus) and direct inhibition of RNase L enzymatic activity (e.g., Theiler’s virus)(Whelan *et al*, 2019; Drappier & Michiels, 2015; Sorgeloos *et al*, 2013). Human immunodeficiency, Influenza A, and vaccinia viruses circumvent this pathway by structurally sequestering dsRNA detection within viral proteins(Chang *et al*, 1992; Schröder *et al*, 1990). Thus, RSV uses a unique mechanism to avoid the OAS-RNase L pathway without halting its function.

RSV dsRNA, the initial trigger of the OAS-RNase L pathway, does not allow for the probing of RSV-infected cells via IFA using anti-dsRNA antibodies; however, it was successfully detected in the purified total RNA (Fig 3, Fig EV4). In RSV-infected cells, IB formation is characteristically observed as a droplet shape, which is enriched with viral proteins such as N, P, M2-1, and L that are linked to replication and transcription(Norrby *et al*, 1970; Wileman, 2007; Carromeu *et al*, 2007; García-Barreno *et al*, 1996; Santangelo & Bao, 2007). These IBs serve as the main sites of new viral RNA synthesis(Rincheval *et al*, 2017). Moreover, RSV facilitates immune evasion by trapping immune factors, such as MDA5, MAVS, and NF-κB subunit p65, in the IBs(Lifland *et al*, 2012; Jobe *et al*, 2020). The dense structural properties of IBs hinder antibodies from accessing internal epitopes, leading to the restriction of fluorescent signals to the IB periphery, complicating the exploration of their internal organization(Rincheval *et al*, 2017). Thus, we hypothesized that dsRNA resides in IBs that act as a barrier to its detection using OASs and anti-dsRNA antibodies. Proteinase K treatment to degrade IBs enabled the detection of dsRNA co-localized with IBs. Furthermore, we disrupted IBs by deleting viral genes necessary for their formation to investigate the ability of IBs to conceal dsRNA and their effect on activating the OAS-RNase L pathway. However, this approach proved unfeasible because the absence of N and P proteins or impaired IB formation critically abrogated viral replication.

Recent studies have identified IBs generated during RSV infection as biomolecular condensates formed via LLPS, similar to those observed in other negative-stranded RNA viruses(Galloux *et al*, 2020; Zhang *et al*, 2024). Suppression of RSV replication after LLPS hardening by A3E or cyclopamine treatment underscores LLPS’s crucial role in RSV replication(Risso-Ballester *et al*, 2021). Surprisingly, treatment with 1,6-HD, which dissolves various types of phase-separated condensates in cells, disrupted the condensation of IBs and induced dsRNA leakage into the cytosol. These results show that RSV may use LLPS to sequester its dsRNA and evade sensing from OASs. Moreover, dsRNA masking by LLPS caused the failure of ectopically activated RNase L in degrading viral RNA. Contrary to our expectations, OAS-RNase L pathway activation was not observed in response to 1,6-HD-induced dsRNA leakage during RSV infection, possibly because of the pleiotropic effects of 1,6-HD. Düster et al. reported that 1,6-HD is unsuitable for investigating the functional relationship between LLPS and cellular pathways(Düster *et al*, 2021; Meduri *et al*, 2022). These results suggest that 1,6-HD impairs the activities of enzymes, including those of kinases and phosphatases, which are crucial for cellular signaling and function. The mechanism of action of 1,6-HD suggests that it exerts non-negligible effects on multiple pathways, including the OAS-RNase L pathway, as demonstrated in our study, although their hypothesis mainly focused on the enzymatic activity of kinases and phosphatases.

In summary, the OAS-RNase L pathway was not activated during RSV infection, and ectopically activated RNase L failed to suppress viral replication. The initial trigger of this pathway produced during RSV replication, dsRNA, was strictly sequestered within the LLPS-mediated IBs but liberated upon LLPS disruption. Notably, transfection with naked dsRNA isolated from RSV-infected cells, which eliminated the protective effect of LLPS, activated the OAS-RNase L pathway. Thus, RSV may evade the OAS-RNase L pathway by sequestering viral RNA via LLPS-driven IB formation (Fig 7). These findings suggest that modulating the physical properties of LLPS structures, such as IBs, could represent a novel antiviral therapeutic strategy against RSV. In addition to hardening IBs(Risso-Ballester *et al*, 2021), approaches aimed at specifically dissolving IBs to expose sequestered viral RNA to the immune system could be explored for their therapeutic potential. This study is the first to demonstrate that LLPS acts as a shielding barrier that protects RNA from RNase-induced degradation. It is unlikely that this evasion strategy is restricted to the OAS-RNase L pathway. Further research is required to determine whether LLPS enables evasion by other immune sensors that detect viral RNA.

**Figure 7-.**
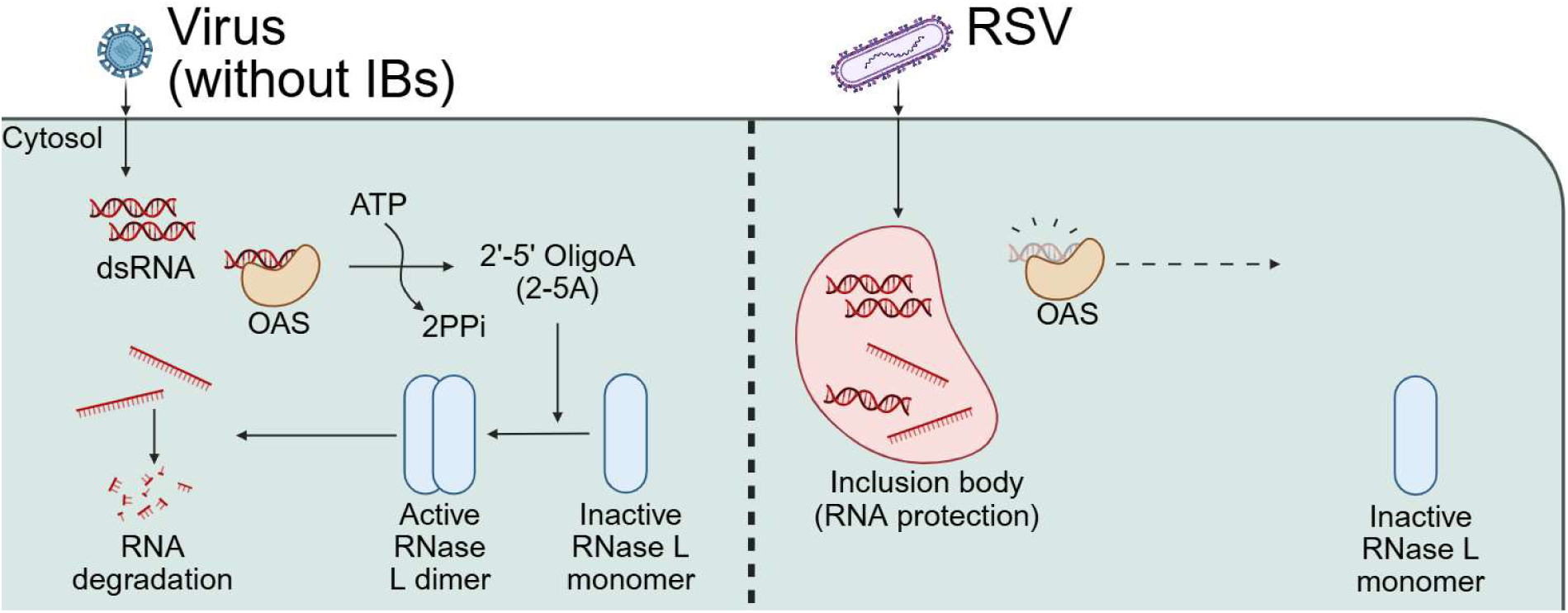
Schematic model of respiratory syncytial virus (RSV) evasion from the dsRNA-mediated oligoadenylate synthetase (OAS)-RNase L pathway. (left) Upon viral dsRNA detection produced by diverse viruses, OASs bind to them and produce 2-5A, which activate RNase L. Cellular and viral single-strand RNAs are degraded, leading to restriction of virus replication and host cell apoptosis. (right) In RSV-infected cells, inclusion bodies (IBs) sequester replication-derived dsRNA, preventing OAS detection and pathway activation. Additionally, IBs shield the viral genome from apically activated RNase L.

## Materials and methods

### Cells and viruses

HEp-2 cells (10023, KCLB) were cultured in minimum essential medium supplemented with 100 U/mL penicillin–streptomycin (Gibco) and 10% (v/v) fetal bovine serum (FBS). A549 cells (ATCC CCL-185) were obtained from the American Type Culture Collection (ATCC; Manassas, VA, USA) and maintained in Dulbecco’s modified Eagle’s medium supplemented with 100 U/mL penicillin– streptomycin and 10% FBS. RSV strains A2 (VR-1540) and 18537 (VR-1580) were acquired from ATCC and amplified within the HEp-2 cells. Both viral strains were propagated three times before use in the experiments. Virus titers were determined via a plaque assay on the HEp-2 cells at 37°C using a methylcellulose overlay for 7 d after infection.

### rRNA cleavage assay

A549 and HEp-2 cells were infected with the indicated virus at an MOI ranging from 0.1–5 or transfected with poly(I:C) (Sigma). Cells were harvested using Trizol reagent (Ambion) at the indicated time points following infection or transfection. rRNA cleavage was analyzed using a 4150 TapeStation System (Agilent Technologies, California, USA) with an RNA ScreenTape (5067-5576, Agilent Technologies).

### Immunoblotting

Cells were lysed in radioimmunoprecipitation assay (RIPA) buffer (RC2002-050-00, Biosesang) for immunoblotting. Proteins were separated via electrophoresis on 10% denaturing polyacrylamide gels and transferred onto polyvinylidene difluoride membranes (Merck Millipore). The membranes were blocked with 5% skim milk (BD Biosciences, Franklin Lakes, NJ, USA) in Tris-buffered saline containing 0.1% Tween 20 for 1 h at 24°C. Subsequently, the membranes were incubated overnight with the following primary antibodies: anti-OAS1 (14498S, Cell Signaling Technology, 1:200), anti-OAS2 (54155S, Cell Signaling Technology, 1:500), anti-OAS3 (21915-1-AP-20, Proteintech, 1:250), anti-RNase L (27281S, Cell Signaling Technology, 1:1,000), anti-β-actin (sc-47778, Santa Cruz Biotechnology, 1:5,000), anti-RSV N (GTX636648, GeneTex, 1:1,000), and anti-ZIKV NS3 (GTX133309, GeneTex, 1:1,000). Proteins were detected using horseradish peroxidase (HRP)-conjugated secondary antibodies (Bio-Rad) and an enhanced chemiluminescence (ECL) reagent (Thermo Fisher Scientific).

RNA dot blot assays to detect dsRNA in cell lysates involved isolating total RNA using Trizol reagent (Invitrogen) following the manufacturer’s instructions, spotting onto nylon membranes (Sigma), and incubating overnight with 5% skim milk (BD Biosciences) in phosphate-buffered saline containing 0.05% Tween 20 (PBS-T) at 4°C. dsRNA was probed using the anti-dsRNA antibodies J2 (76651, Cell Signaling Technology, 1:1,000) and 9D5 (Ab00458-23, Absolute antibody, 1:1,000) for 2 h at 24°C. For dsRNA visualization, HRP-conjugated secondary antibodies (Bio-Rad) and ECL reagents (Thermo Fisher Scientific) were used.

### Quantitative reverse transcription polymerase chain reaction (RT-PCR)

RT-qPCR (QuantStudio 3, Applied Biosystems, Foster City, CA, USA) was performed using One-Step PrimeScript III RT-qPCR Mix (Takara Bio, Shiga, Japan). The RSV F gene was detected using a probe-based qPCR assay (Integrated DNA Technologies, Coralville, IA, USA). Additionally, the *IFN-β*, *OAS1*, *OAS2*, and *OAS3* genes were detected using individual customized probes (Integrated DNA Technologies). Supplementary Table S1 lists the sequences of the qPCR probes and primers used in this study.

### Immunofluorescence staining

A549 cells were seeded onto 8-chamber slides and cultured overnight. Cells were infected with RSV A2 at an MOI of 2, fixed with 4% paraformaldehyde for 20 min at the indicated time points post-infection, permeabilized using 0.1% Triton X-100 in PBS-T for 20 min, and blocked with 5% bovine serum albumin for 1 h at 24°C. The slides were incubated overnight with the following primary antibodies: anti-dsRNA antibodies J2 and 9D5, anti-RSV N, anti-RSV F (01-07-0121, Cambridge, 1:1,000), and anti-RSV P (ab94965, Abcam, 1:1,000). Subsequently, the slides were rinsed three times with PBS-T and incubated with the appropriate Alexa Fluor-conjugated secondary antibodies (Invitrogen) for 1 h at 24°C. For nuclear staining, 4’,6-diamidino-2-phenylindole (DAPI) was used.

To disrupt the LLPS, 5% 1,6-HD dissolved in MEM was prepared, and the culture medium was replaced with this solution at 12 or 24 h post-infection for 0 to 30 min. After treatment, the medium containing 1,6-HD was removed, and the cells were rinsed three times with PBS. Immunofluorescence staining was performed as previously described.

### Naked dsRNA purification from RSV-infected cells

Naked dsRNA from RSV-infected cells was isolated using a previously reported protocol, with some modifications(Decker *et al*, 2019). Briefly, HEp-2 cells were infected with RSV A2 at an MOI = 2. After 48 h, the total RNA was extracted using Trizol reagent. For dsRNA enrichment, anti-dsRNA monoclonal antibody J2 pre-conjugated to protein A/G magnetic beads (20424, Thermo) was incubated with total RNA in a pre-chilled reaction buffer (20 mM Tris-HCl [pH 7.0], 150 mM NaCl, 0.2 mM EDTA, 0.2% Tween-20) at 4°C for 2 h with rotational mixing. Using a microspin column (89879, Thermo), beads were recovered and washed three times with wash buffer (20 mM Tris-HCl [pH 7.0], 150 mM NaCl, 0.1 mM EDTA, 0.1% Tween-20). After resuspending the beads in 150 μL of wash buffer, bound dsRNA was eluted, followed by the addition of 450 μL Trizol LS reagent (10296010, Invitrogen). Finally, the dsRNA was isolated according to the manufacturer’s instructions.

### Statistical analysis

All experiments were conducted at least three times. All data were analyzed using GraphPad Prism 8.0 (GraphPad Software, San Diego, CA, USA). Statistical significance was set at P < 0.05. The figure legends include the description of the specific analytical methods.

## Acknowledgments

This research was conducted under Project No. KK2432-20 (A Study on the Convergence Platform for Infectious Disease Diagnosis and Prevention). Young-Chan Kwon is supported by the Korea Research Institute of Chemical Technology (KRICT) and by a National Research Foundation of Korea (NRF) grant funded by the Ministry of Education, Science, and Technology (MIST) of the Korean government RS-2023-00208568). Soohwan Oh is supported by a Basic Science Research Program through the National Research Foundation of Korea (NRF), funded by the Ministry of Education (2022R1C1C1010699) and the Ministry of Science and ICT (MSIT), Korea, under the Information Technology Research Center (ITRC) support program (IITP-2025-RS-2023-00258971), supervised by the Institute for Information & Communications Technology Planning & Evaluation (IITP).

## Disclosure and competing interest statment

The authors declare no conflicts of interest.

---

**Expanded View Figure 1-.**
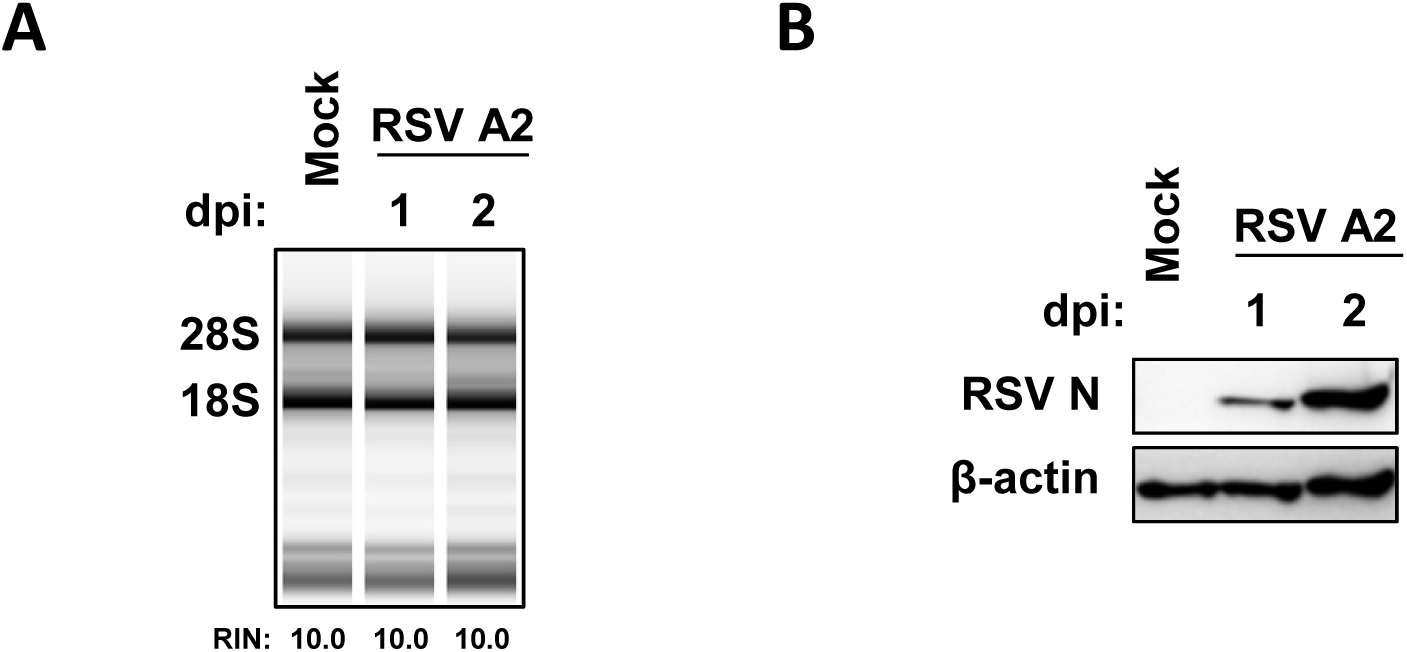
RSV infection does not induce OAS-RNase L activation in the HEp-2 cells. (A, B) HEp-2 cells were infected with RSV A2 (MOI = 2) and lysed at the indicated time points. rRNA cleavage and RNA integrity number (RIN) were analyzed using the RNA TapeStation System. Viral replication was examined by immunoblotting with anti-RSV N and anti-β-actin antibodies.

**Expanded View Figure 2-.**
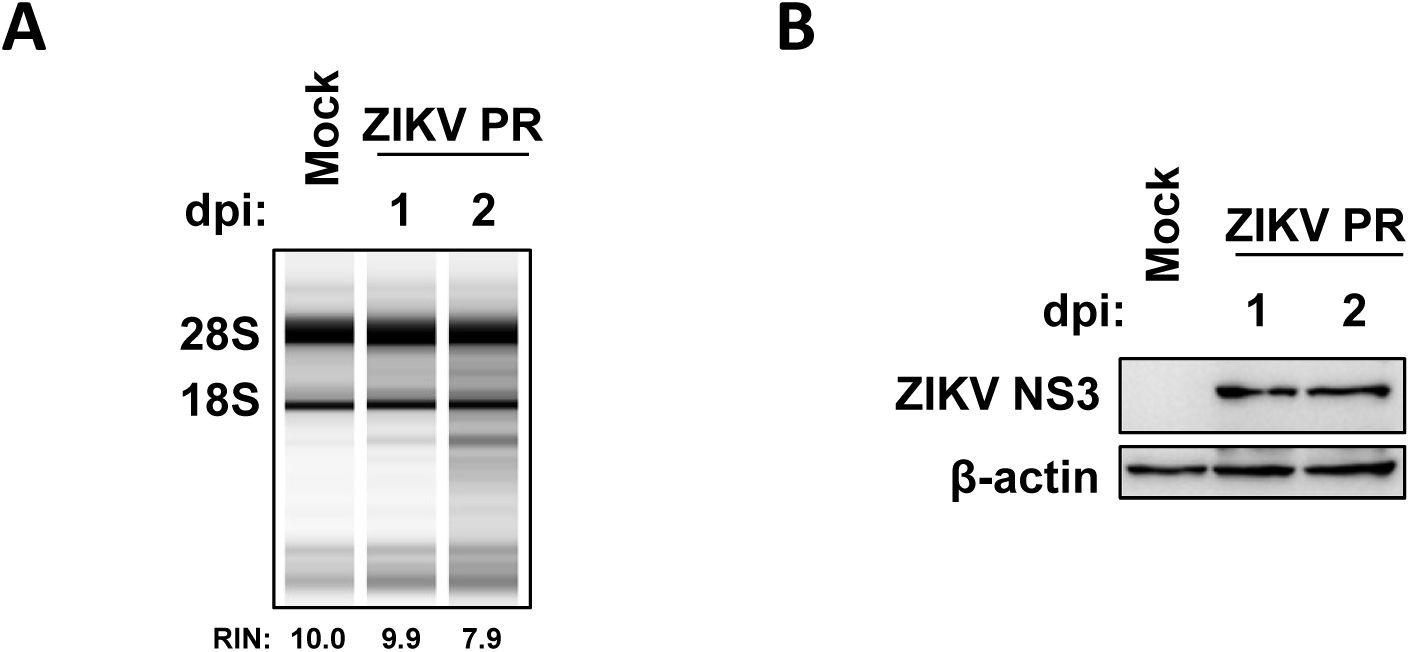
ZIKV infection activates the OAS-RNase L pathway. (A, B) The A549 cells were infected with ZIKV PRVABC59 (multiplicity of infection [MOI] = 2). RNA and protein were harvested and examined using the RNA TapeStation System and immunoblotting with anti-ZIKV NS3 and anti-β-actin antibodies.

**Expanded View Figure 3-.**
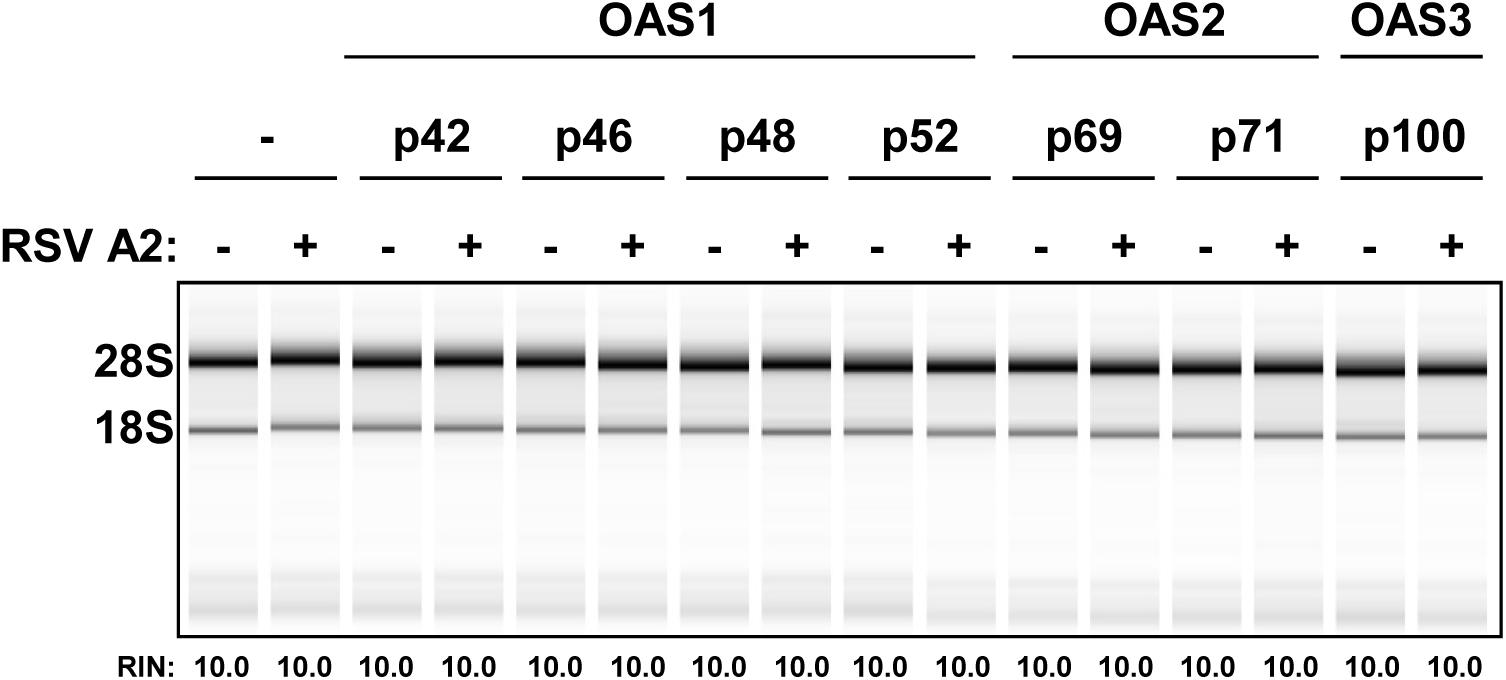
RNase L was not activated in the RSV-infected A549 cells overexpressing OAS isoforms. The A549 cells were transfected with the OASs isoforms one day before RSV infection (MOI = 2). Two days after infection, the RNA was purified and analyzed using the RNA TapeStation System.

**Expanded View Figure 4-.**
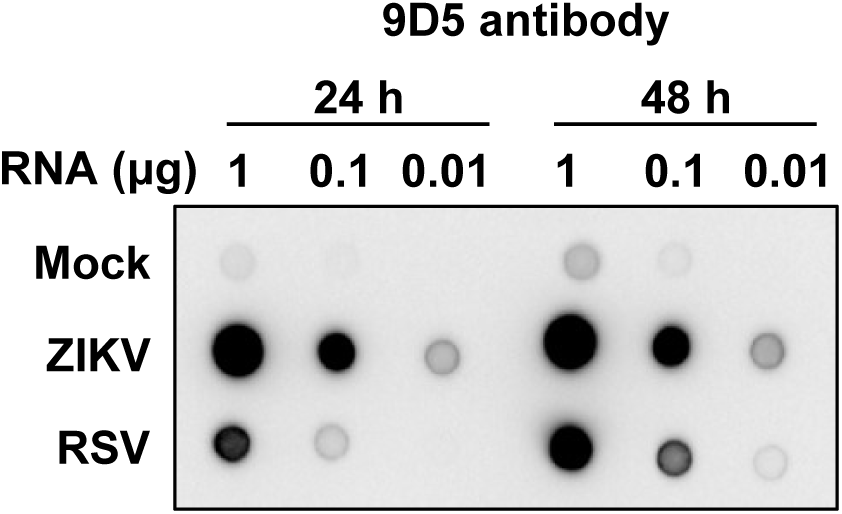
The dsRNA detection in total RNA extracted from the RSV-infected cells. Total RNA from the mock-, RSV-, and ZIKV-infected cells was transferred onto nylon membranes. The dsRNA was detected using an anti-dsRNA antibody (9D5).

**Expanded View Figure 5-.**
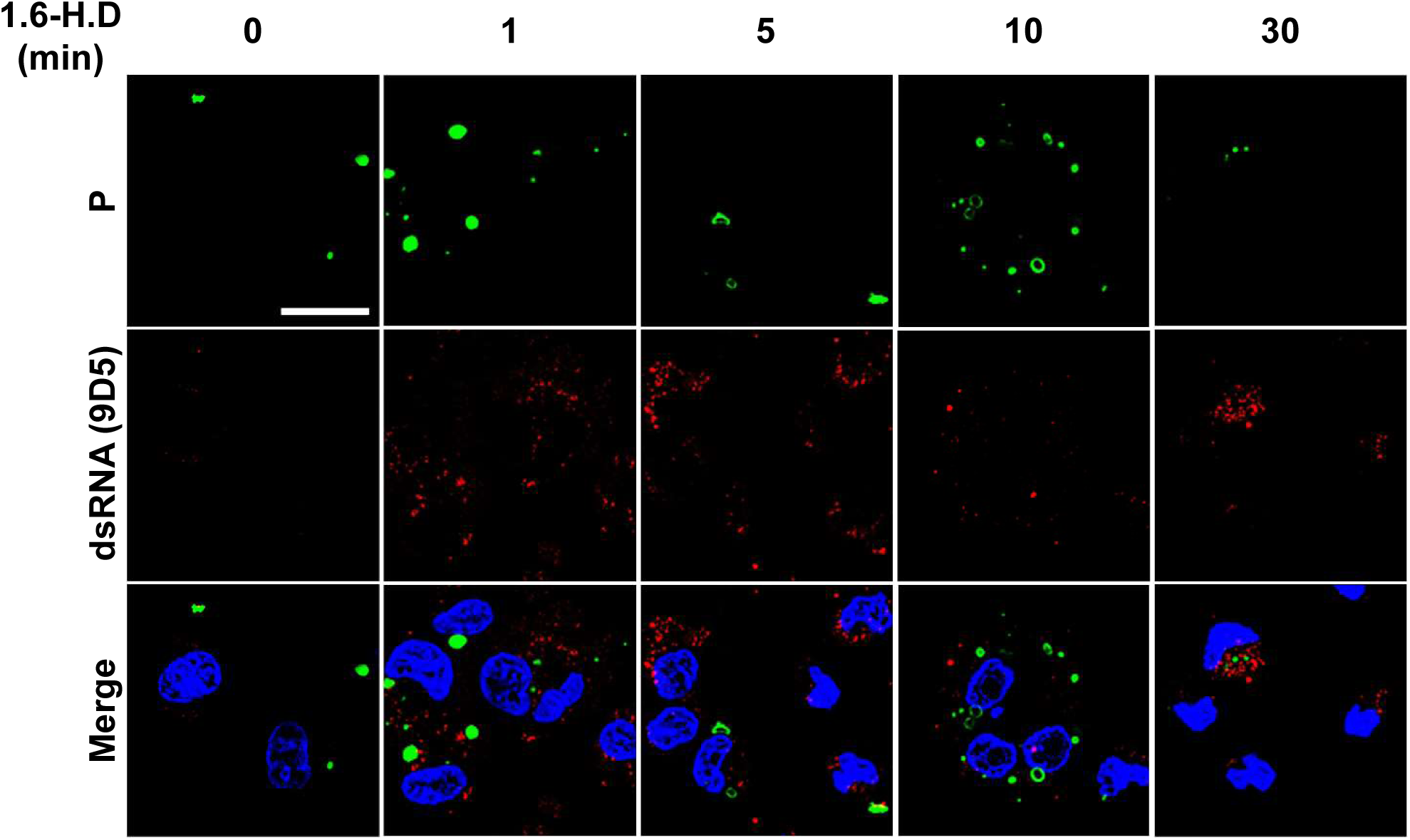
Disruption of IBs with an LLPS inhibitor induces dsRNA leakage into the RSV-infected cells. The A549 cells were infected with RSV A2 (MOI, 2). Before the cells were fixed 24 h after infection, they were treated with 5% 1,6-HD at the indicated times. Cells were stained with anti-P antibody (green) and anti-dsRNA (9D5, red). Scale bar: 20 μm.

**Expanded View Figure 6-.**
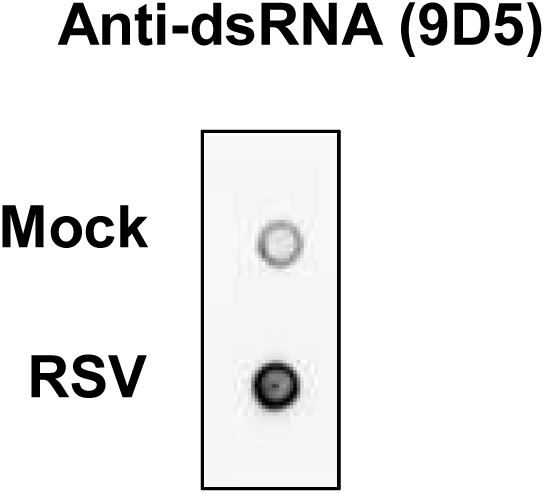
Naked dsRNA purified from the RSV-infected cells. After dsRNA purification, dsRNA enrichment was confirmed by dot blotting using an anti-dsRNA antibody (9D5).

**Expanded View Figure 7-.**
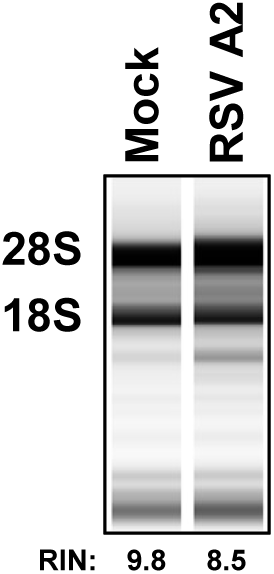
Transfection of the total RNA extracted from the RSV-infected cells activates the OAS-RNase L pathway. The total RNA was purified from mock- and RSV-infected cells and transfected into A549 cell with 10 μg. At 24 h after transfection, the RNA was assessed using the RNA TapeStation System.

**Expanded View Table - 1 qRT-PCR primers and probes.**

